# Leveraging heterogeneity across multiple data sets increases accuracy of cell-mixture deconvolution and reduces biological and technical biases

**DOI:** 10.1101/206466

**Authors:** Francesco Vallania, Andrew Tam, Shane Lofgren, Steven Schaffert, Tej D Azad, Erika Bongen, Meia Alsup, Michael Alonso, Mark Davis, Edgar Engleman, Purvesh Khatri

## Abstract

*In silico* quantification of cell proportions from mixed-cell transcriptomics data (deconvolution) requires a reference expression matrix, called basis matrix. We hypothesized that matrices created using only healthy samples from a single microarray platform would introduce biological and technical biases in deconvolution. We show presence of such biases in two existing matrices, IRIS and LM22, irrespective of the deconvolution method used. Here, we present immunoStates, a basis matrix built using 6160 samples with different disease states across 42 microarray platforms. We found that immunoStates significantly reduced biological and technical biases. We further show that cellular proportion estimates using immunoStates are consistently more correlated with measured proportions than IRIS and LM22, across all methods. Importantly, we found that different methods have virtually no effect once the basis matrix is chosen. Our results demonstrate the need and importance of incorporating biological and technical heterogeneity in a basis matrix for achieving consistently high accuracy.

## Introduction

Cell-mixture deconvolution is an established *in silico* approach for quantifying cell subpopulations directly from bulk gene expression data of mixed cell samples^1–4^. Multiple computational methods have been developed over the years^5,6^ to estimate the proportions of immune cells in blood^6^ and tissue biopsies^7^, as well as cell-type specific expression profiles from bulk expression data^8,9^. The underlying assumption in virtually every deconvolution approach to date is that the observed expression of any given gene in a mixed-tissue sample is a combination of its expression across each cellular subset^1^. Based on this assumption, methods for estimating cellular frequencies from mixed tissue data use a variant of regression model, such as linear regression^3^, quadratic programming^4^, robust regression^7^, or support vector regression^7^. Irrespective of the type of statistical model, each method requires a reference expression matrix, called a basis matrix, that is composed of genes specifically expressed in the expected cell subsets found in the tissue of interest^8^.

Typically, a basis matrix is constructed from sorted cell expression data by combining expression profiles of sorted immune cells from one (e.g., IRIS^23^) or more datasets (e.g., LM22^7^), which are profiled using a single microarray platform to ensure homogeneity in expression data. However, it is possible that this approach can introduce a technical bias in a basis matrix towards the microarray platform used for transcriptome profiling, resulting in lower deconvolution accuracy for samples that are profiled using different platforms. Furthermore, basis matrices to date have been created using expression data solely from healthy subjects^2,3,7^, which can further introduce biological bias that could affect deconvolution accuracy and limit their applicability to disease samples.

Here, we show that current deconvolution approaches are significantly affected by severe technical and biological bias by measuring accuracy across 5479 human transcriptomes. We found that the presence of these biases significantly reduced deconvolution accuracy. We therefore hypothesized that a basis matrix created by integrating data from multiple independent cohorts of healthy and disease samples profiled using different microarray platforms would reduce biological and technical biases and improve accuracy of deconvolution.

To test this hypothesis, we created a new basis matrix, immunoStates, that leverages biological and technical heterogeneity across 6,160 whole transcriptomes of human sorted blood cells measured on 42 microarray platforms. We used a multi-cohort analysis framework that leverages biological and technical heterogeneity present across multiple independent studies for this purpose. The approach has been previously shown to increase reproducibility in gene expression signatures across a broad spectrum of diseases including organ transplant, sepsis, infectious diseases, cancer, and systemic sclerosis^10–20^. We show that immunoStates allows for accurate deconvolution with no significant bias across different microarray platforms and disease samples. Importantly, our analyses showed that, for any given basis matrix, different deconvolution methods produce highly correlated results, demonstrating that the choice of the matrix is more important than the deconvolution method itself. Our findings provide strong evidence for the importance of the basis matrix in determining deconvolution accuracy. We conclude that incorporation of technical and biological heterogeneity in the construction of the matrix reduces bias, which increases accuracy in cell mixture deconvolution independently of the method.

## Results

### Using a single microarray platform introduces technical bias in a basis matrix

We hypothesized that a basis matrix created using gene expression data generated from a single microarray platform would exhibit significant platform-dependent bias (technical bias) in deconvolution accuracy. To test our hypothesis, we used IRIS and LM22 to deconvolve 17 independent datasets consisting of 1010 whole transcriptome profiles of human peripheral blood mononuclear cells (PBMCs) measured across eight microarray platforms from two different manufacturers (see Methods). Both basis matrices were constructed using only healthy samples profiled only on Affymetrix microarrays^3,7^. Further, to generalize our findings across multiple methods, we used four deconvolution algorithms (linear regression, quadratic programming, robust regression, and support vector regression)^3,4,7^. We estimated accuracy of deconvolution across samples by computing their “goodness of fit” as previously described (see Methods)^7^. Briefly, if the original mixed-tissue sample expression could be reconstituted by combining the estimated proportions of individual cell types with the expression from basis matrix, we would observe high goodness of fit and expect good deconvolution accuracy.

We observed significant differences in goodness of fit between microarray platforms across both matrices, irrespective of the method used (**Figure 1A-B**), demo nstrating the presence of platform-specific bias. We quantified the extent of these differences for each basis matrix using median absolute deviation (MAD) of goodness of fit, a measure of heterogeneity robust to outliers as described before^21^, across samples from different platforms. We calculated MAD as difference in goodness of fit for each sample from mean goodness of fit across all platforms for a given basis matrix, and estimated its statistical significance against the null hypothesis that there was no technical variation between samples (see Methods). We observed significant heterogeneity in goodness of fit between platforms for both IRIS (MAD = 0.18, p = 2.27e-9) and LM22 (MAD = 0.12, p = 4.79e-4), irrespective of the method used (**Supplementary Figure 1**).

**Figure 1:**
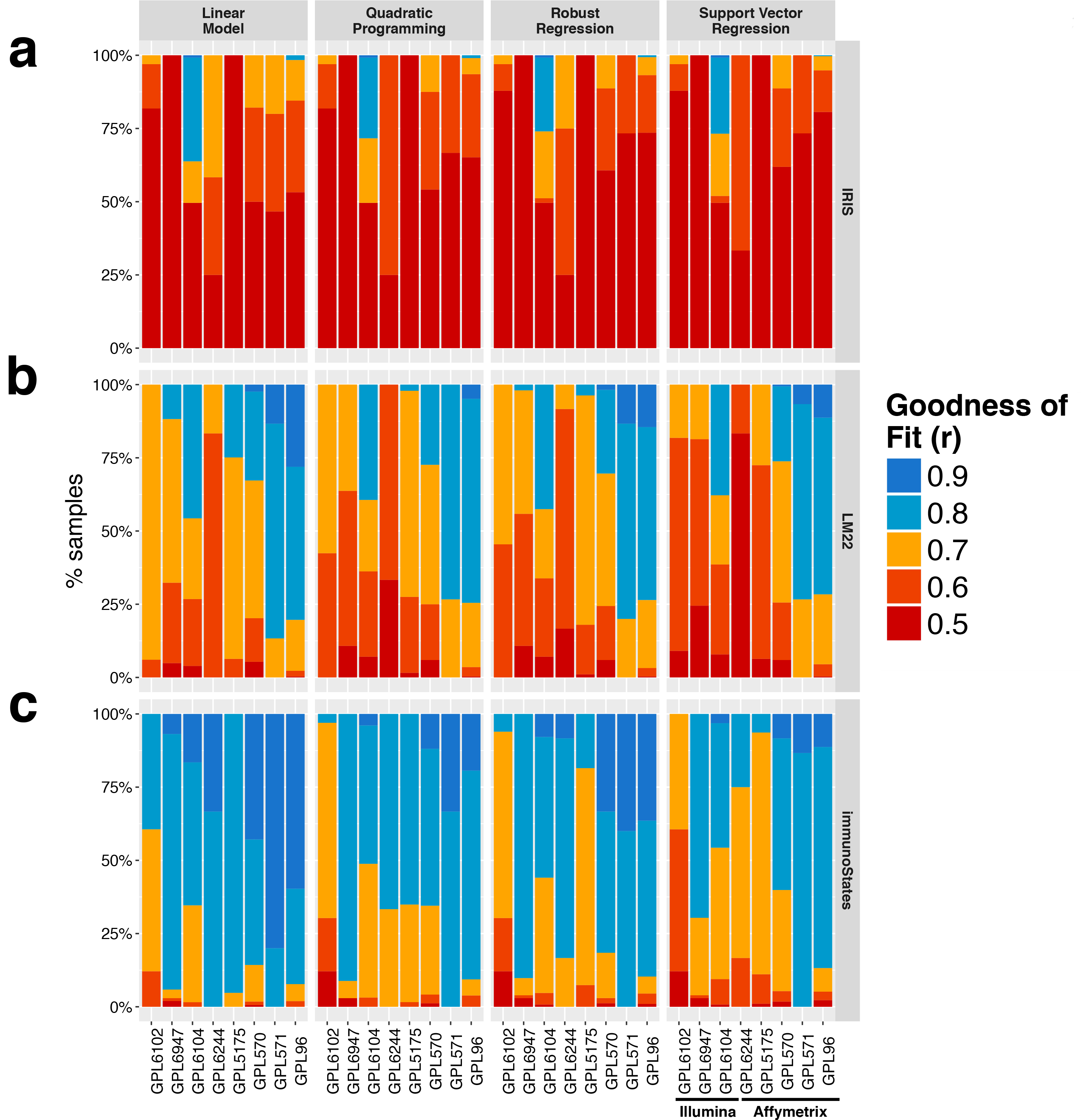
Analysis of platform bias in deconvolution across multiple methods and signature matrices. **(a)** Goodness of fit values across 1010 human PBMC samples as a function of microarray platform using the IRIS signature matrix. Goodness of fit is displayed as a stacked barplot with color indicating corresponding values starting from goodness of fit value of 0.5 or lower up to values of 0.9 and above. Barplots are grouped by the method of deconvolution used for the analysis. **(b)** Same as in **(a)** for LM22. **(c)** Same as in **(a)** for immunoStates.

### Leveraging heterogeneity across microarray platforms reduces platform-dependent technical bias in a basis matrix

Next, we hypothesized that a basis matrix created using multiple microarray platforms will reduce platform-dependent technical bias in cellular deconvolution. We collected 165 publicly available gene expression datasets from GEO that profiled 6160 samples from 20 sorted human blood cell types using 42 microarray platforms (**Supplementary Figure 2A**, see Methods). We did not discard experiments based on sorting strategy, platform manufacturer, or disease state of the sample.

Using these data, we created a new basis matrix consisting of 317 cell-type specific genes, called immunoStates, (**Supplementary Figure 2B**, see Methods), of which only 12 genes were shared between IRIS, LM22, and immunoStates (**Supplementary Figure 2C**). A large fraction of genes in immunoStates (76%) was not shared with IRIS or LM22. We then deconvolved the technical bias evaluation cohort using immunoStates as a basis matrix across all four methods. Unlike IRIS and LM22, we observed no heterogeneity in goodness of fit between microarray platforms (**Figure 1C**; MAD = 0.05, p = 0.20; **Supplementary Figure 1**). Importantly, mean goodness of fit using immunoStates was significantly higher than IRIS (p < 2.2 e-16) and LM22 (p < 2.2 e-16) irrespective of the method used. Together, our results demonstrate that a basis matrix created using heterogeneous data from multiple platforms reduces technical bias.

### Leveraging heterogeneity across healthy and disease samples reduces biological bias in a basis matrix

Cell quantification technologies, such as FACS and CyTOF, use a set of predefined phenotypic markers that are not affected by either disease-or treatment-induced changes^22^. For instance, irrespective of whether a sample is from a healthy control or a patient with or without any treatment, CD14 and CD56 are used to identify monocytes and natural killer cells, respectively. Similarly, a basis matrix should be unaffected by disease-and treatment-induced changes to be broadly applicable across a large number of diseases and conditions.

We hypothesized that a basis matrix created using only healthy samples (e.g., IRIS and LM22) will have lower goodness of fit when deconvolving a disease sample, and hence, lower deconvolution accuracy, whereas a basis matrix created using both healthy and disease samples (e.g., immunoStates) will have higher goodness of fit and accuracy. To test this hypothesis, we used E-MTAB-62, a gene expression compendium of 5372 samples representing primary tissues and cell lines^23^. For the purpose of this analysis, we considered only primary samples from human subjects, consisting of 4,067 blood-and tissue-derived samples from either healthy donors or individuals with a disease, such as leukemia, solid tumors, and neurodegenerative disorders. For each pair-wise combination of a basis matrix and a deconvolution method, we determined the effect of disease on deconvolution accuracy by estimating its ability to distinguish blood-from tissue-derived samples based on significance of goodness of fit, as described previously^7^.

First, because each of the three matrices only contained blood cells, we would expect them to have a significantly higher goodness of fit values for blood-derived samples compared to those for solid tissue samples as they are not represented in any of the basis matrices. In 1,383 healthy samples, irrespective of the method, the goodness of fit was higher for blood-derived samples than tissue-derived samples for all basis matrices (**Supplementary Figure 3A**). This result translated into an accurate distinction of blood from tissue-derived samples for all matrices and methods based on significance of deconvolution (AUCs: IRIS 93.60 ± 0.01%; LM22 95.61 ± 0.01%; immunoStates 93.75 ± 0.01%, **Figure 2A**) for healthy samples.

**Figure 2:**
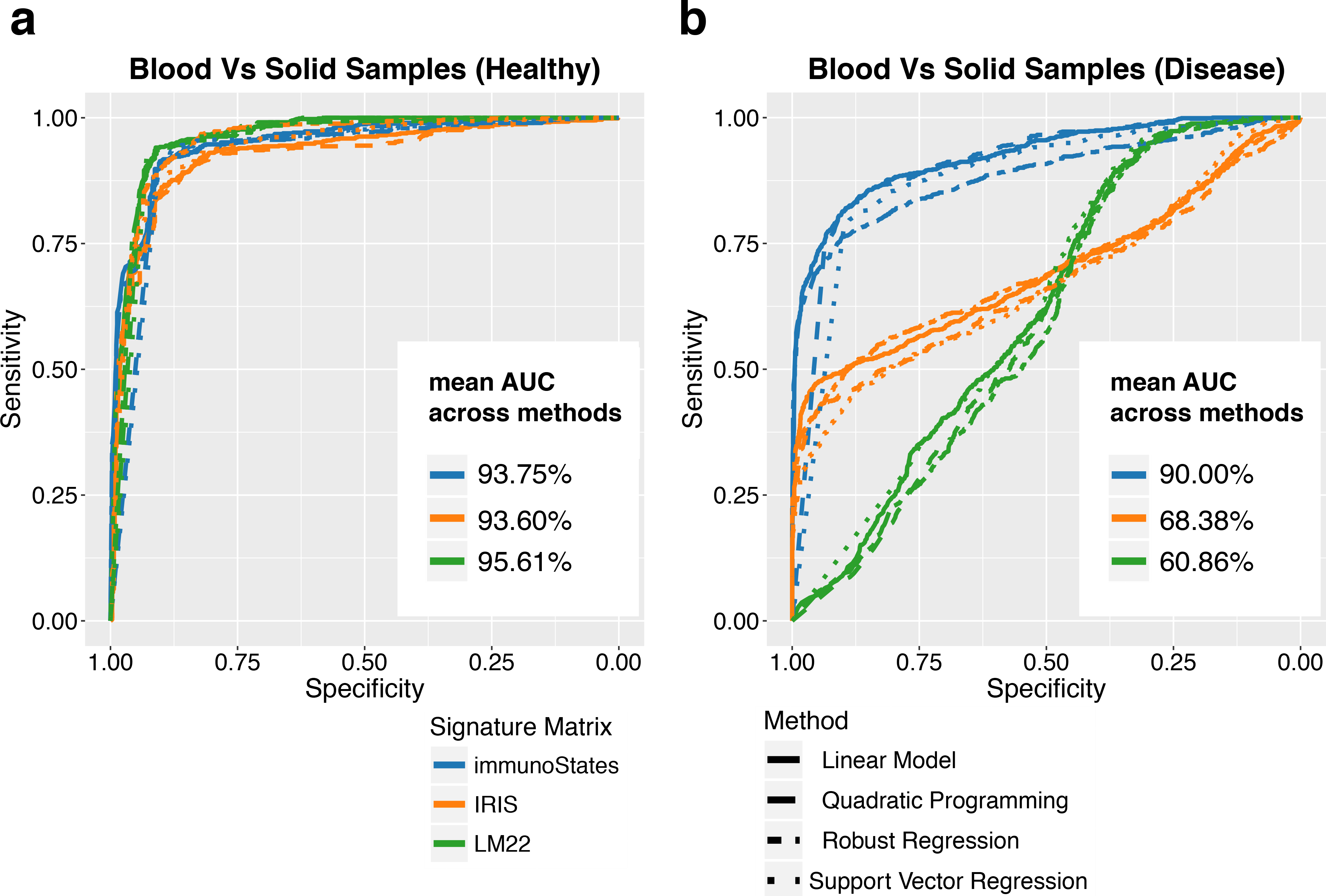
Effect of disease on deconvolution. **(a)** ROC curves indicating the ability of IRIS, LM22, and immunoStates (denoted by line color) to distinguish blood-derived samples from tissue biopsies in healthy donors (1383 samples) using goodness of fit across all tested methods (denoted by line type). AUCs indicate mean AUC for an individual signature matrix across all methods **(b)** Same as in **(a)** but in disease samples (2684 samples).

Second, if the expression of genes in a basis matrix changed in a disease sample, we would expect low goodness of fit for both blood and solid tissue samples, indicating lower deconvolution accuracy. For 2,684 disease samples, when using IRIS or LM22, the goodness of fit values for blood-and tissue-derived samples were highly similar (**Supplementary Figure 3A**) and resulted in lower discrimination between them (IRIS: AUC = 68.38 ± 0.01% and LM22: AUC = 60.86 ± 0.01%) (Figure 2B). In contrast, immunoStates had significantly higher goodness of fit for blood-derived samples than tissue-derived samples, irrespective of the deconvolution method used, resulting in high accuracy for distinguishing blood-and tissue-derived disease samples (AUC = 90.00 ± 0.01%, **Figure 2B**, **Supplementary Figure 3B**). Collectively, these results demonstrated that a basis matrix created using only healthy transcriptome profiles contains a biological bias against disease samples, which makes it difficult to distinguish between blood and solid tissue samples, and results in lower deconvolution accuracy. In contrast, creating a basis matrix using both healthy and disease samples significantly reduce the biological bias and increase deconvolution accuracy.

### Different methods produce highly correlated cellular proportions for a given basis matrix

Our results revealed that incorporating biological and technical heterogeneity by using both healthy and disease samples profiled across multiple platforms in a basis matrix reduced platform and disease bias irrespective of the deconvolution method used. Therefore, we tested whether the basis matrix had a stronger effect on the deconvolution results than the method used to estimate cell proportions. Interestingly, for a given basis matrix, all methods produced high correlated cell proportion estimates (r = 0.777 ± 0.013 **Figure 3**). In contrast, for a given method, we observed significantly lower correlations in cell proportion estimates when using different basis matrices (r = 0.409 ± 0.022, P < 2.2e-16), or when both matrix and method were different (r = 0.392 ± 0.013, P < 2.2e-16). We observed these trends irrespective of whether the sample came from blood or solid tissue biopsies (**Supplementary Figure 4**). These results provide a strong evidence that the basis matrix is the major determinant of deconvolution accuracy, not the method used for deconvolution, and demonstrate that virtually no method can overcome biological and technical bias present in a basis matrix.

**Figure 3:**
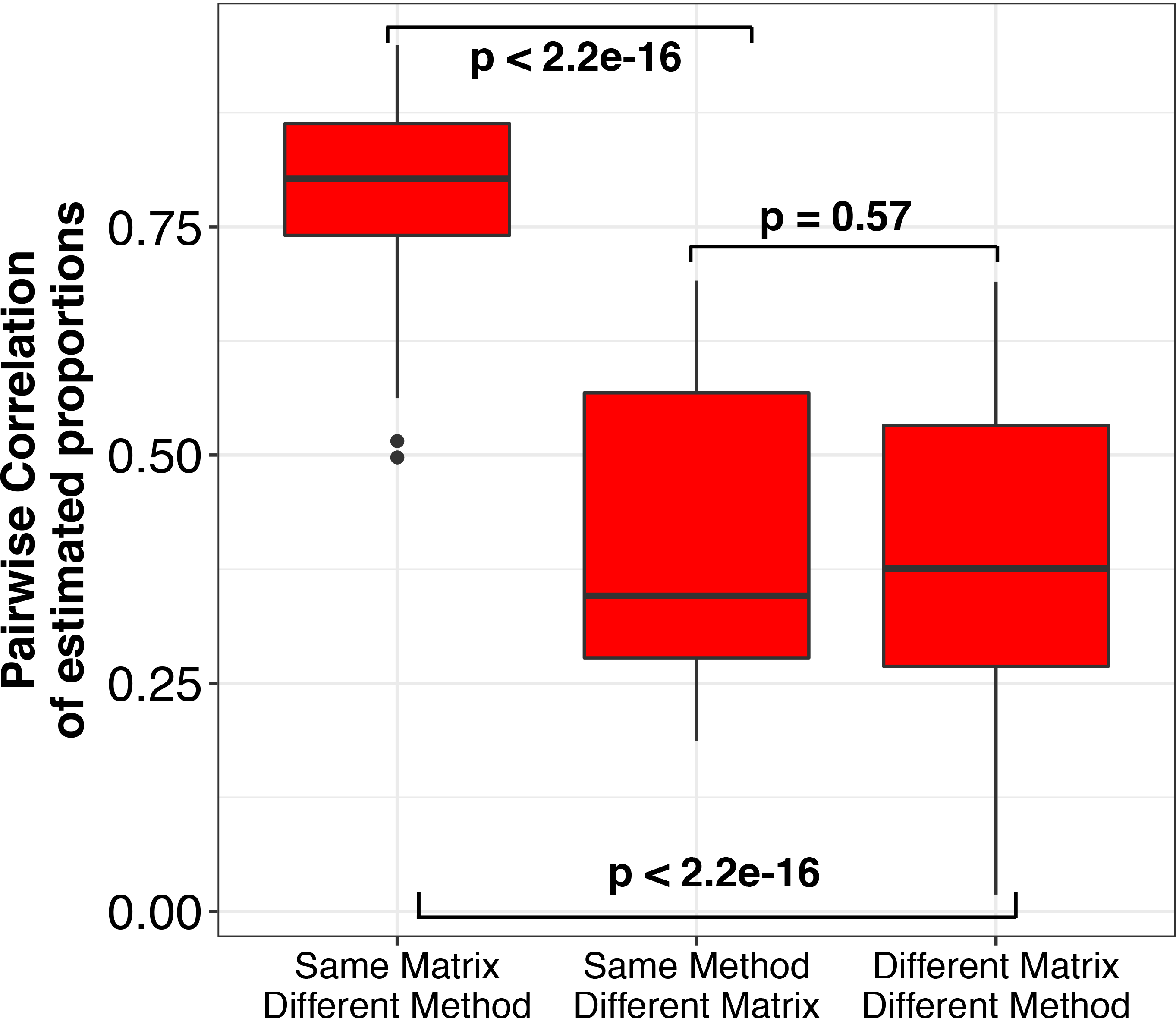
Deconvolution concordance by signature matrix and method. Boxplots representing the distribution of pairwise correlation coefficients between estimated proportions for all matrices and deconvolution methods. Comparisons were divided in (1) pairs with the same signature matrix but run with different methods, (2) pairs with different signature matrices but run using the same method, and (3) pairs where both matrix and method were different. Significance analysis was performed using the Wilcoxon’s paired rank sum test.

### Reduction of technical and biological bias in a basis matrix increases accuracy of deconvolution

Next, we explored whether the reduction biological and technical bias in a basis matrix results in increased accuracy of deconvolution by correlating estimated cell proportions with cell proportions measured using FACS or Coulter counter for each pair of a basis matrix and a deconvolution method. We identified five gene expression datasets of 402 human whole blood or PBMC samples with paired cell counts data available (see Methods). These datasets were generated using Illumina HT12 V4.0 or Affymetrix Primeview microarrays as follows: (1) two independent datasets consisting of 176 healthy human PBMC samples profiled using Illumina HT-12 V4.0 arrays paired with flow-cytometry data (GSE65133, GSE59654)^7,24^, and (2) a whole blood dataset of 226 healthy samples, a subset of which were profiled over three consecutive years using Affymetrix PrimeView arrays (see Methods).

Across the five datasets, estimated cell proportion by IRIS and LM22 had significantly lower correlations with measured cell proportions (IRIS: r = 0.15 ± 0.07, p = 1.9e-5; LM22: r = 0.56 ± 0.06, p = 5.9e-4) compared to immunoStates (r = 0.80 ± 0.03) (**Figure 4A**, **Supplementary Figures 5-9**). In concordance with previous report, we found that IRIS and LM22 systematically over- or underestimated individual cellular proportions even for high frequency cell subsets such as Monocytes and CD4+ T-cells (**Supplementary Figure 10**)^7^. Importantly, across all basis matrices, no method produced consistently more correlations with measured cell proportions (Linear Model p = 0.41; Quadratic Programming p = 0.49; Robust Regression p = 0.46; Support Vector Regression p = 0.55), and provided further evidence that the accuracy of deconvolution is determined by a basis matrix instead of a deconvolution method.

**Figure 4:**
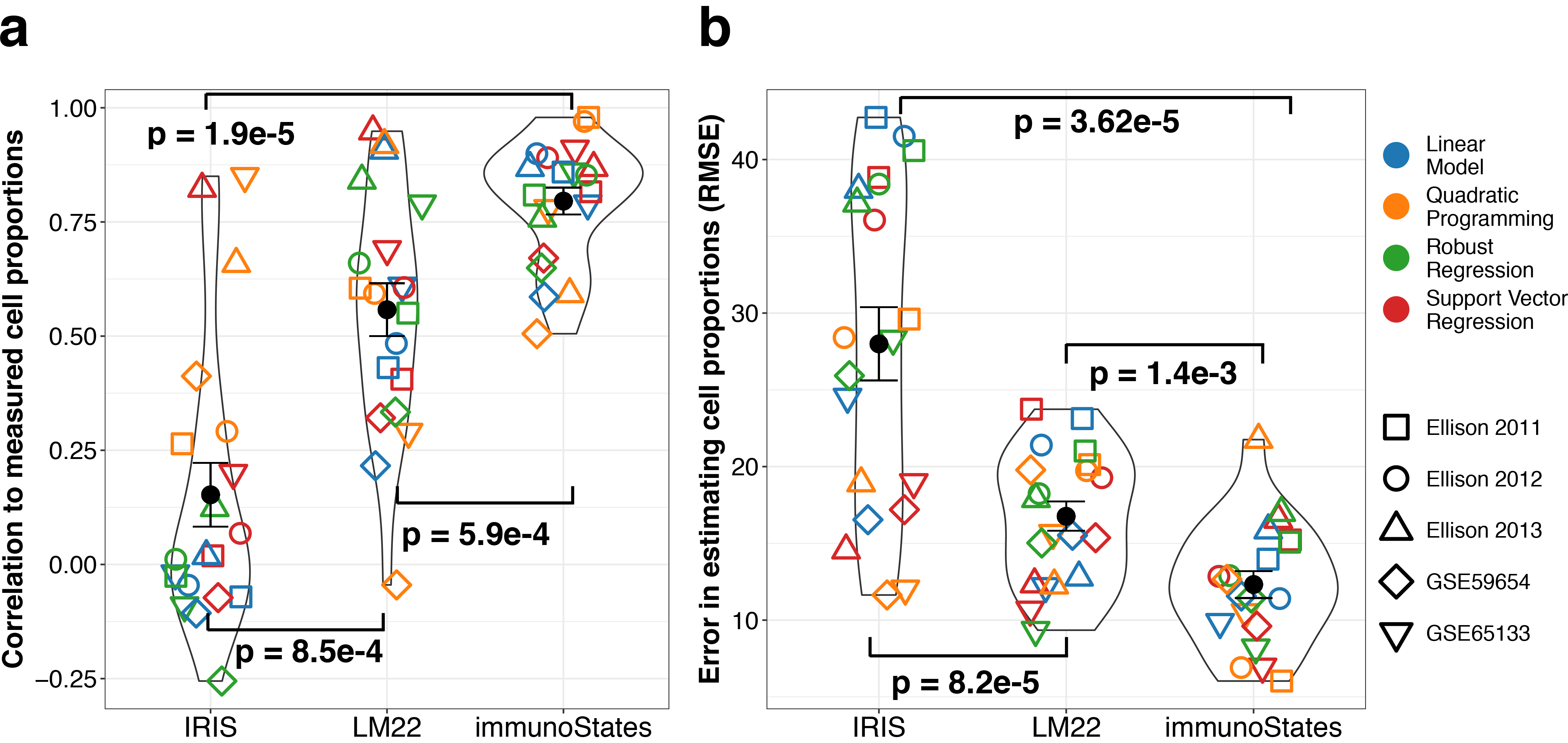
Comparison between estimated and measured cell proportions across 402 human blood samples. **(a)** Correlation between measured cell proportions and deconvolution estimates in five different human sample cohorts (denoted by different shapes) across different deconvolution methods (denoted by different colors) using IRIS, LM22, and immunoStates (x-axis). Correlation is measured by Pearson’s correlation coefficient. **(b)** Same as in **(a)** for RMSE between measured and estimated cell proportions.

Finally, we quantified the extent of over- and under-estimation of cell proportions across all cohorts and methods by computing the root mean square error (RMSE) between measured and estimated proportions (**Figure 4B**). In comparison to IRIS (RMSE = 27.99 ± 2.39, p = 3.6e-5) and LM22 (RMSE = 16.78 ± 0.95, p = 1.4e-3), we found that that immunoStates generated estimates with significantly lower RMSE (RMSE = 11.93 ± 0.88) across all cohorts and deconvolution methods used. Again, no single method had significantly lower RMSE than others across all matrices and cohorts (Linear Model p = 0.62; Quadratic Programming p = 0.49; Robust Regression p = 0.40; Support Vector Regression p = 0.52). Collectively, our results provide strong evidence that accuracy of current deconvolution methods is affected by biological and technical biases present in a basis matrix, and no method is able to overcome these biases. Our results further demonstrate that creating a basis matrix by leveraging biological and technical heterogeneity across multiple independent cohorts reduces these biases, and significantly improves accuracy of deconvolution irrespective of the statistical model used.

## Discussion

Using whole transcriptome profiles, cell-mixture deconvolution methods quantify cell subsets within a mixed-tissue sample without physical separation of its components. We hypothesized that the current practice of creating a basis matrix for deconvolution using only healthy samples profiled using the same microarray platform introduces biological and technical biases that reduces accuracy of deconvolution. Our analysis of two basis matrices, IRIS and LM22, both created using only one microarray platform and healthy samples, showed significant heterogeneity in deconvolution results between different microarray platforms, and lower discriminatory power for distinguishing blood and solid tissue samples when obtained from a patient instead of a healthy control.

There is increased evidence that leveraging biological and technical heterogeneity across multiple independent datasets identifies robust and reproducible disease signatures^10–20^. Here, we hypothesized that the heterogeneity present in publicly available datasets can be used to create a basis matrix with significantly reduced biological and technical bias, and increase accuracy of deconvolution. We used 165 publicly-available gene expression datasets that profiled 6,160 sorted human immune cell samples using 42 different microarray platforms to create a 317-gene basis matrix called immunoStates. Our analysis showed that immunoStates substantially reduced technical and biological bias and resulted in more accurate cell proportion estimates.

Unexpectedly, we found that the accuracy of all basis matrices was independent of the statistical method used for deconvolution. For a given basis matrix, all methods produced highly correlated cellular proportions. This result has important long-term implications as virtually all efforts to date have been focused on developing new methods to improve deconvolution accuracy rather than the design of the basis matrix^6^. This result also suggests that immunoStates may be readily applied to any future deconvolution strategy as long as they rely on the use of a basis matrix to estimate cell proportions.

Our approach has an important limitation. We leveraged heterogeneity present in public data to create immunoStates. Despite more than 1 million human transcriptome profiles in NCBI GEO and EBI ArrayExpress, more specific and rare cell subsets are less likely to have sufficient heterogeneous data across multiple platforms. It is not clear how much data is “sufficient” to represent cellular heterogeneity. Our previous work has suggested that 4-5 datasets consisting of approximately 250 samples may be enough to represent disease heterogeneity^17^. However, the heterogeneity at cellular level may require more datasets. We expect that continued accumulation of additional sorted-cell datasets in public repositories over time will increase both the accuracy and the breadth of future basis matrices. We also encourage researchers profiling sorted cells to make them available publicly. Second, arguably, all basis matrices require *a priori* knowledge of the populations within the sample of interest. However, current cell sorting methods also require *a priori* selection of markers. The advantage of immunoStates is that through increased accuracy and reduced technical, biological, and methodological bias, it can leverage existing data in public repositories to identify cellular subsets that should be further explored using targeted technologies such as FACS or CyTOF. By avoiding cellular subsets that are not changing in existing data and avoiding further profiling them, immunoStates can help researchers design better experiments that increase the probability of identifying relevant and novel cell subsets.

## Material and Methods

### Dataset collection and pre-processing

Unless otherwise noted, we downloaded all datasets from Gene Expression Omnibus (GEO, www.ncbi.nlm.nih.gov/geo/) using the MetaIntegrator package from CRAN^19^. We then normalized each expression data set using quantile normalization. We computed gene-level expression for each sample by averaging expression values from probes mapping to the same genes while excluding individual probes that were promiscuously associated to more than one transcript. Datasets used to estimate technical bias are described in Supplementary Table 1. Dataset E-MTAB-62 was downloaded, processed, and annotated from Array Express (http://www.ebi.ac.uk/arrayexpress) using the ArrayExpress R package. Dataset GSE65133 and its paired flow-cytometry data were directly downloaded from the CIBERSORT website (https://cibersort.stanford.edu). Paired flow-cytometry data for GSE59654 was downloaded from ImmPort (https://immport.niaid.nih.gov/; study ID: SDY404). Data from the Stanford-Ellison cohort was collected and processed as previously described^25^. Datasets used to estimate accuracy by comparing with measured cell counts are described in Supplementary Table 2. Dataset E-MTAB-62 was downloaded, processed, and annotated from Array Express (http://www.ebi.ac.uk/arrayexpress) using the ArrayExpress R package.

### Creation of the immunoStates signature matrix

We collected and processed 166 publicly available gene expression datasets comprising 6240 microarray samples profiling sorted human leukocytes. Datasets used to build immunoStates are described in Supplementary Table 3. All datasets were converted in gene-specific expression matrices using the original probe annotation files available from GEO and then combined into a single expression matrix using quantile normalization. Each sample was first annotated following experimental description from the original study, resulting in 47 different cell types. From these initial annotations, cells were grouped into more general categories in order to increase number of studies and platforms represented in each cell type. We defined 20 different cell types: CD4+ T-cells, CD8+ T-cells, gamma-delta T-cells, CD14+ monocytes, CD16+ monocytes, macrophages M0, macrophages M1, macrophages M2, CD56-high natural killer cells, CD56-dim natural killer cells, naïve B-cells, memory B-cells, plasma cells, myeloid dendritic cells, plasmacytoid dendritic cells, hematopoietic progenitors, MAST cells, neutrophils, eosinophils, and basophils. Cell types were then grouped under manually defined “lineages” according to their biological similarity: T-cells, monocytes, macrophages, natural killer cells, B-cells, myeloid dendritic cells, plasmacytoid dendritic cells, hematopoietic progenitors, and granulocytes (see Supplementary Table 3 for annotation details). For each cell type within each lineage, we then computed Hedge′s *g* effect sizes to determine differential expression comparing a given cell type (case) against all remaining cells within that lineage (controls). For each gene ***g*** and cell type ***i*** within lineage ***l***, we then computed a differential effect size defined as 
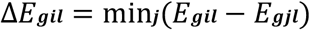
where ***j*** is any cell type within lineage ***l*** such that ***j*** ≠ ***i***. A high differential effect size indicates a strong separation between our target cell type ***i*** and its closest cell type ***j***. We then ranked genes in decreasing order according to their differential effect sizes, and performed a step-wise search to identify the smallest gene signature able to accurately classify cell type ***i*** from every other cell type ***j*** in ***l***. We first estimated the classification accuracy of the first gene of the list by computing a ROC curve and estimating the area under the curve (AUC) and then we incrementally added one gene at a time following their ranking and recomputed the AUC corresponding to the new gene set. We repeated this process until we identified the minimal gene set that produced an AUC proximal to the maximum (with e = 0.005), requiring a minimum of 5 genes per signature. We performed this strategy on each cell type across each lineage and obtained a signature of 214 genes. We then applied the same strategy to distinguished lineages from one another. We excluded all the signature genes that were used to separate cell types due to their confounding contribution in separating lineages. We performed the same gene selection strategy and obtained a second gene set of 98 genes. Together, these sets make the 312 genes that form immunoStates. To build the signature matrix, we computed the mean expression for every gene in every cell type (312 genes by 20 cell types) from the quantile normalized expression matrix. All code was written and run using the R programming language.

### Cell-mixture deconvolution

We performed deconvolution with support vector regression using the CIBERSORT algorithm (v1.03) as described previously^7^. We implemented linear regression, quadratic programming, and robust regression using existing R programs and packages (lm, quadprog, and MASS) as previously described^3,4,7^. We replicated all the pre-processing steps that CIBESORT performed on both the expression sample and basis matrix (quantile normalization and re-scaling of the matrix) in order not to be confounded when we compared different methods. We downloaded the LM22 basis matrix from the CIBERSORT website (https://cibersort.stanford.edu) and the IRIS matrix from the CellMix R package. Statistical comparisons were performed using the Wilcoxon’s rank sum test. Analysis and plots were generated using the R programming language.

### Analysis of platform-dependent technical bias in deconvolution

To quantify the extent of platform-dependent technical bias in deconvolution, we analyzed a collection of 1125 microarray samples profiling human PBMCs in diseased and healthy individuals profiled on platforms GPL96, GPL571, GPL570, GPL5715, GPL6244, GPL6104, GPL6947, GPL6102, and GPL6480 (Supplementary Table 1). We measured deconvolution performance using the goodness of fit score as described previously^7^. Briefly, for a given a basis matrix **M** and a mixture sample gene expression vector s, deconvolution estimates the known cell proportion vector **p** such that 
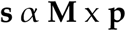

Goodness of fit of the basis matrix **M** is defined as the Pearson′s correlation coefficient between **s** and the reconstituted expression vector **ŝ** defined as **M** × **p**. This value is indicative of how well a particular basis matrix **M** fits **s**. Tests comparing goodness of fits of individual platform/manufacturer were performed for each method using the Wilcoxon’s rank sum test and were then integrated into a final p-value using Fisher’s log sum rule. We calculated heterogeneity for a given matrix by computing the median absolute deviation (MAD) of the median goodness of fit across every platform and deconvolution method. We chose MAD because of its robustness to outliers in estimating the heterogeneity of the distribution of interest^21^. To estimate significance of our MAD scores, we generated a background distribution of MAD scores for each platform, which represent an expected distribution of homogenous MAD values, and then performed a Z-test, asking whether the observed MAD was significantly higher than expected. All analysis and plots were generated using the R programming language.

### Analysis of disease effect to deconvolution

To estimate the effect of disease to sample deconvolution, we analyzed dataset E-MTAB-62 which contains samples profiled on Affymetrix HG-U133A arrays (GPL96). For the purpose of this analysis, we excluded all cell line samples and removed samples that had been used to generate immunoStates. We labeled blood-derived samples as cases (607 and 776 in healthy and disease-affected subjects respectively) and solid-tissue biopsies as controls (934 and 1750 in healthy and disease-affected subjects respectively). We deconvolved each sample, estimated the significance of the goodness of fit as previously described^7^, and used it as a score to distinguish blood-derived samples from biopsies. We measured accuracy of classification by computing a ROC curve and estimating the area under the curve (AUC). Analysis and visualization was performed using the R programming language.

---

**Supplementary Figure 1:**
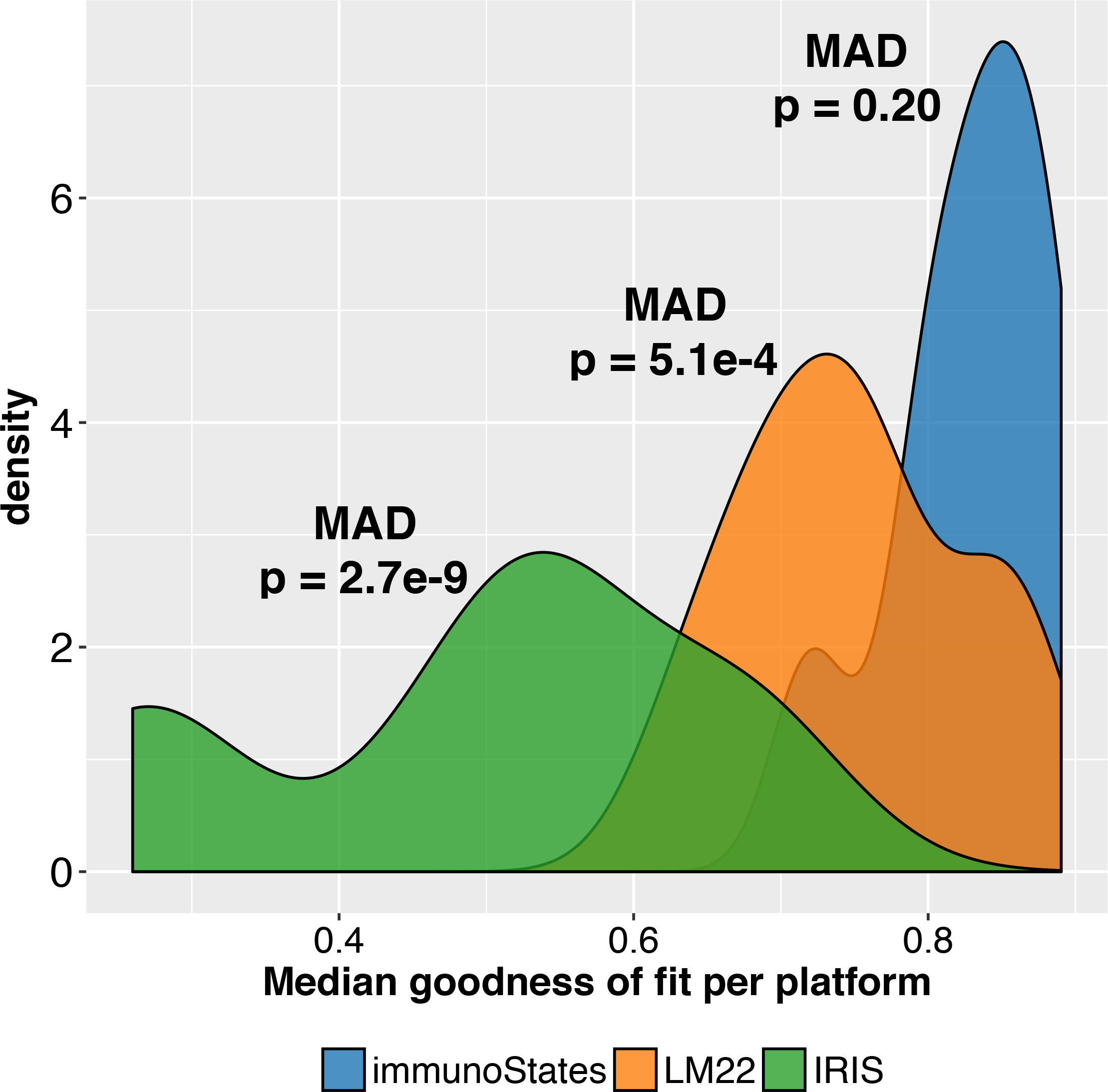
Platform bias in cell mixture deconvolution. Density plots representing the distribution of median goodness of fit for each platform across all methods grouped by different matrices (represented by fill color). Significance of platform bias is computed by estimating the Median Absolute Distance (MAD) of each distribution and comparing it to a null distribution that assumes no technical variation between samples.

**Supplementary Figure 2:**
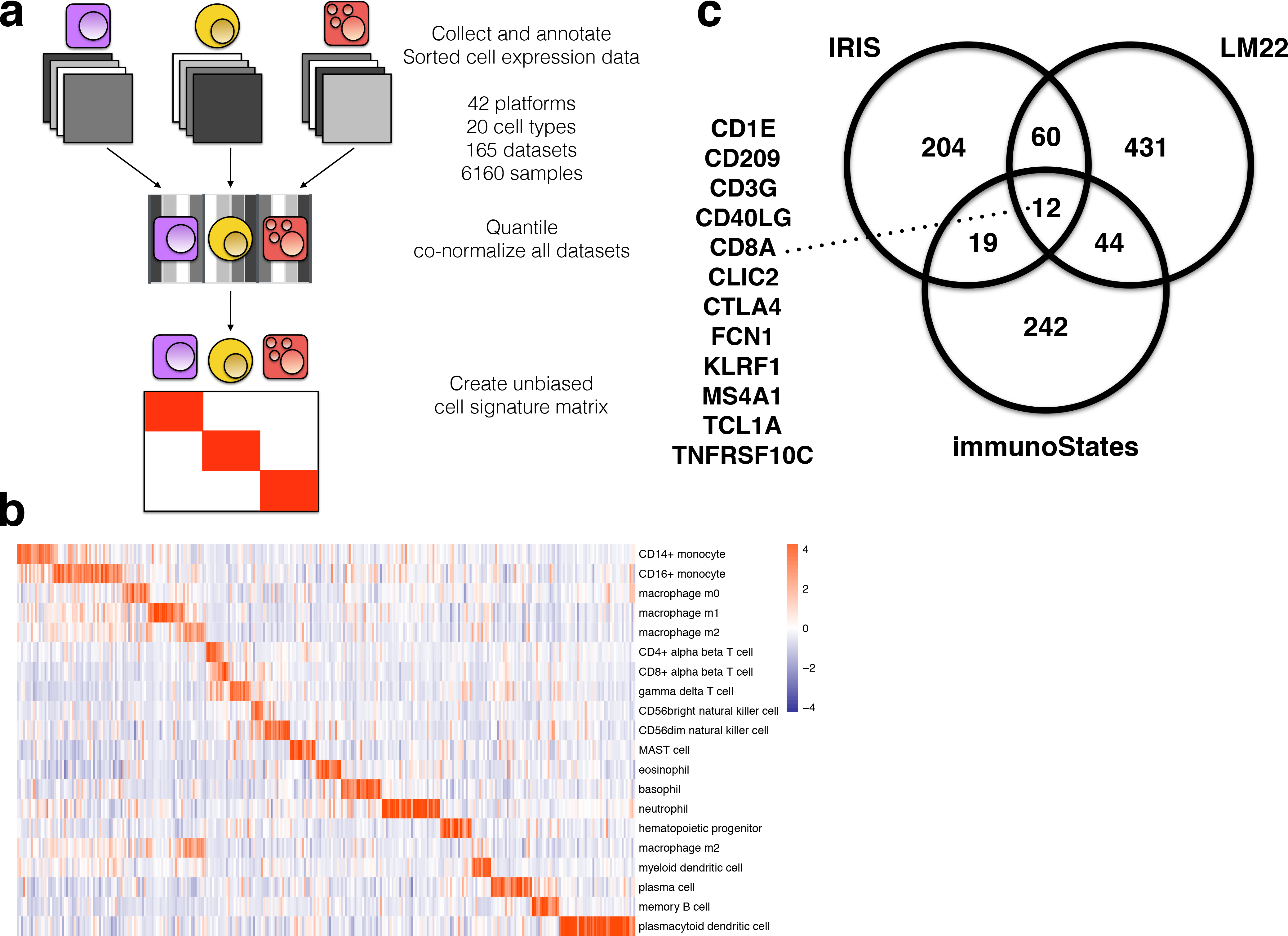
Creation of the immunoStates signature matrix. **(a)** Flow-chart describing the steps for the creation of the immunoStates expression matrix. **(b)** Heatmap showing expression of the immunoStates signature genes in target cell types. Genes expression values are displayed as z-scores per gene across all cell types. **(c)** Venn-diagram depicting the overlap between gene-sets between each basis matrix. Genes overlapping across all three matrices are listed on the left side of the diagram.

**Supplementary Figure 3:**
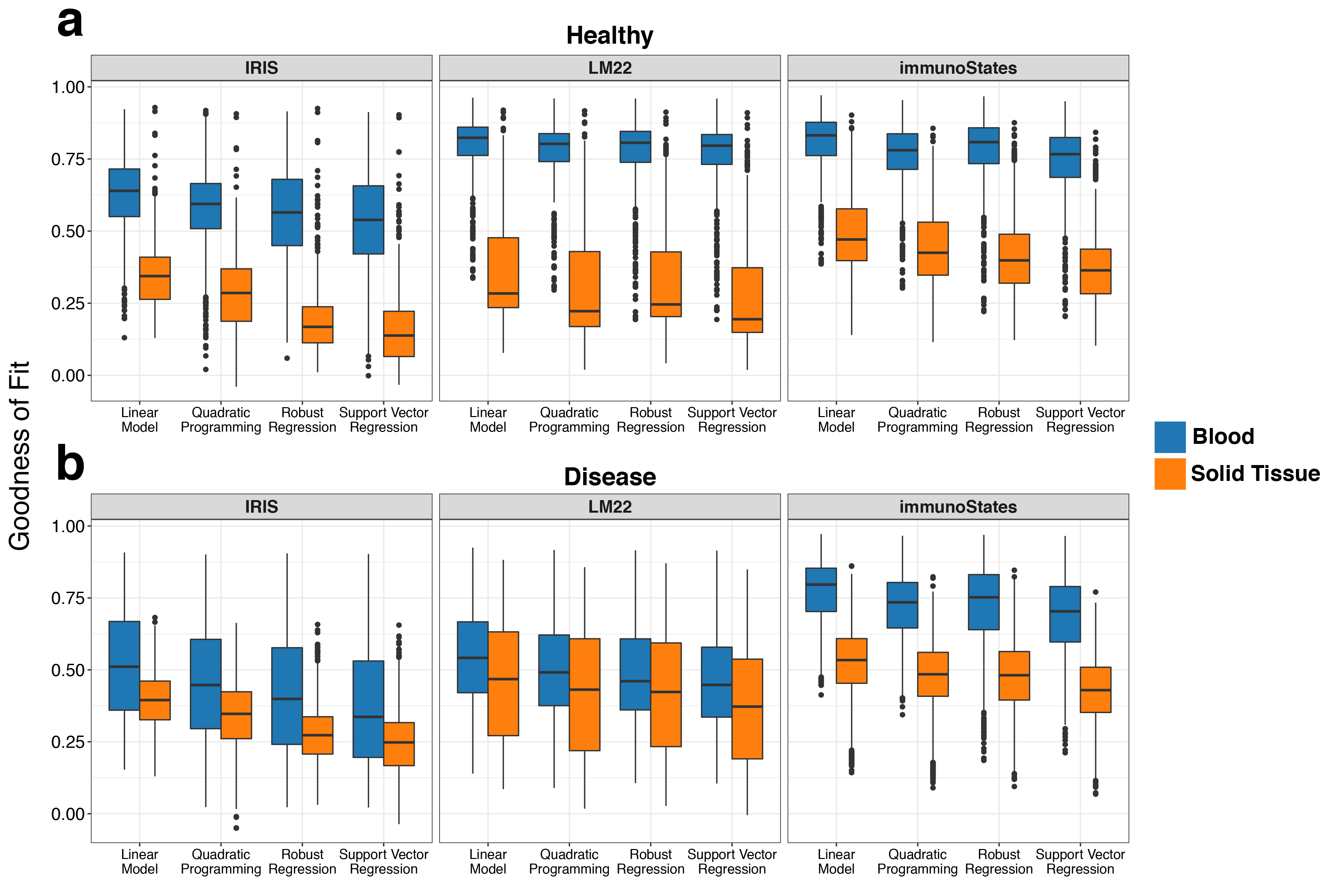
Goodness of fit in healthy and diseased samples. (a) Boxplots indicating goodness of fit scores (y-axis) for blood-derived and tissue-derived samples in healthy donors (1383 samples) across multiple deconvolution methods (x-axis) for IRIS, LM22, and immunoStates. (b) Same as in (a) but in disease samples (2684 samples).

**Supplementary Figure 4:**
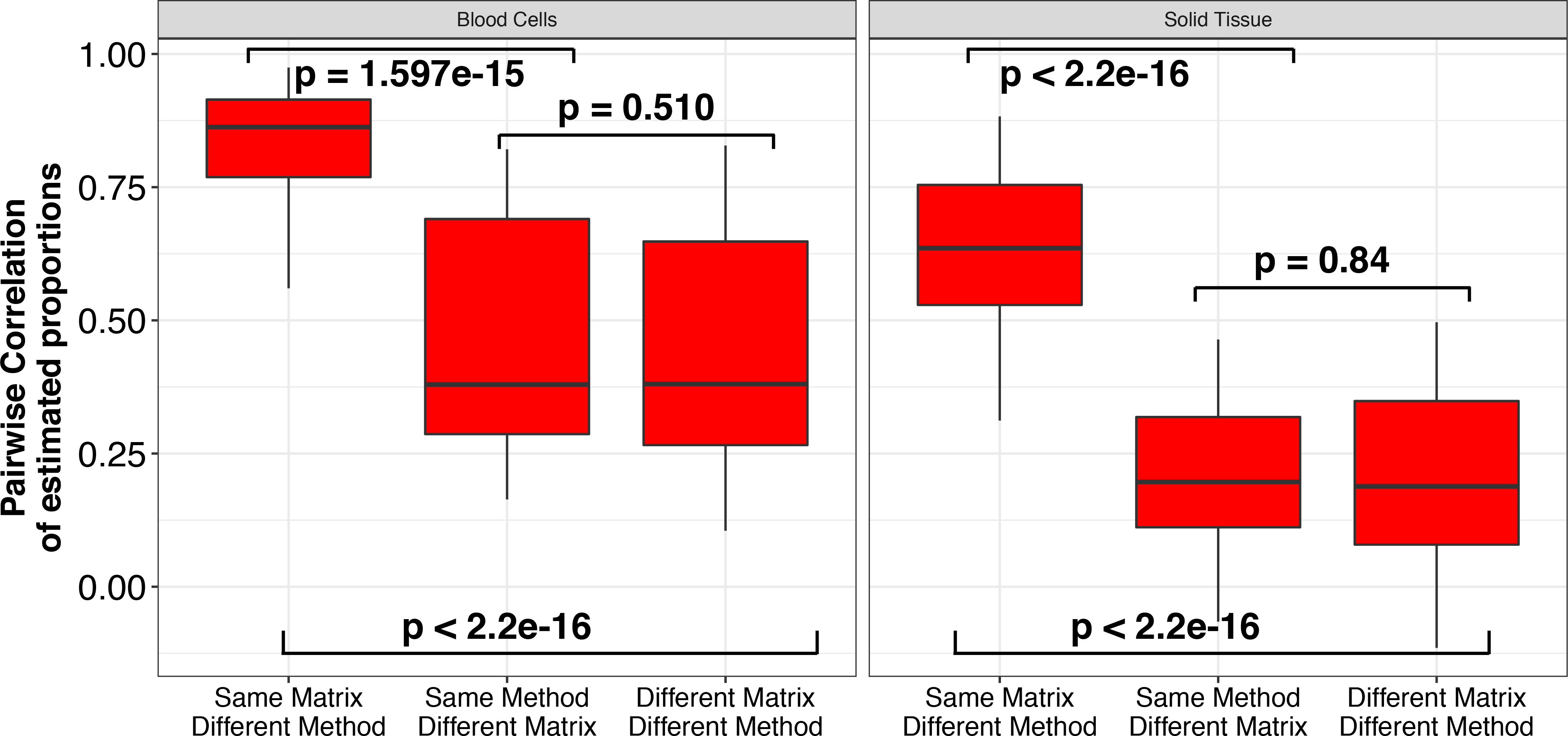
Deconvolution concordance by signature matrix and method across blood and solid tissue. Boxplots representing the distribution of pairwise correlation coefficients between estimated proportions for all matrices and deconvolution methods. Comparisons were divided in (1) pairs with the same signature matrix but run with different methods, (2) pairs with different signature matrices but run using the same method, and (3) pairs where both matrix and method were different. Significance analysis was performed using the Wilcoxon’s paired rank sum test. Results are shown for samples containing blood cells or solid tissue biopsy from Lukk *et al* 2010.

**Supplementary Figure 5:**
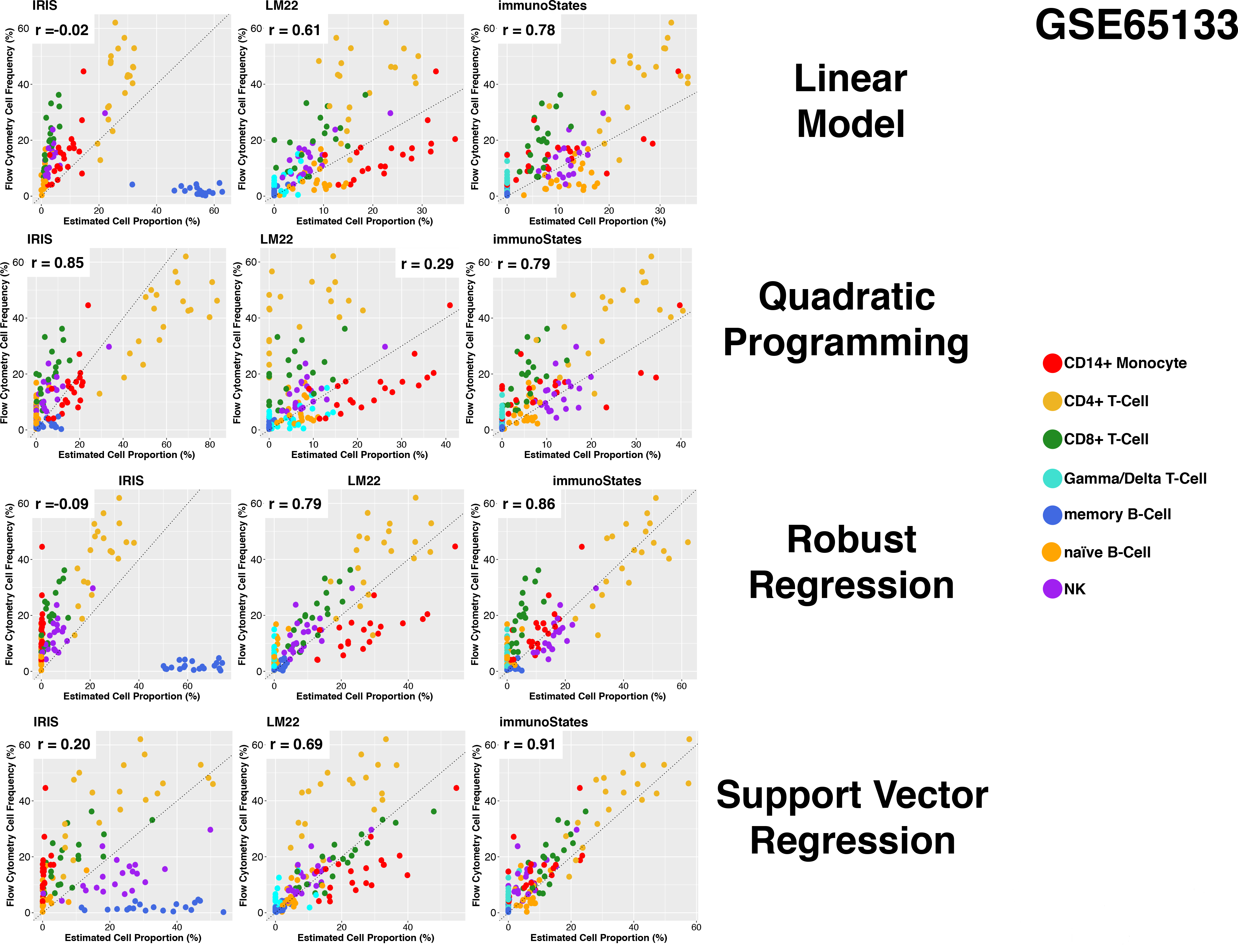
Correlation plots between measured and estimated proportions. **(5)** Correlation plots of estimated (x-axis) and measured cell proportions (y-axis) for each method and matrix combination for samples in GSE65133. Correlation is measured by Pearson’s correlation coefficient. **(6)** Same as in **(5)** for GSE59654. **(7,8,9)** Same as in **(5)** for Stanford-Ellison 2011, 2012, and 2013 sample cohorts respectively.

**Supplementary Figure 6:**
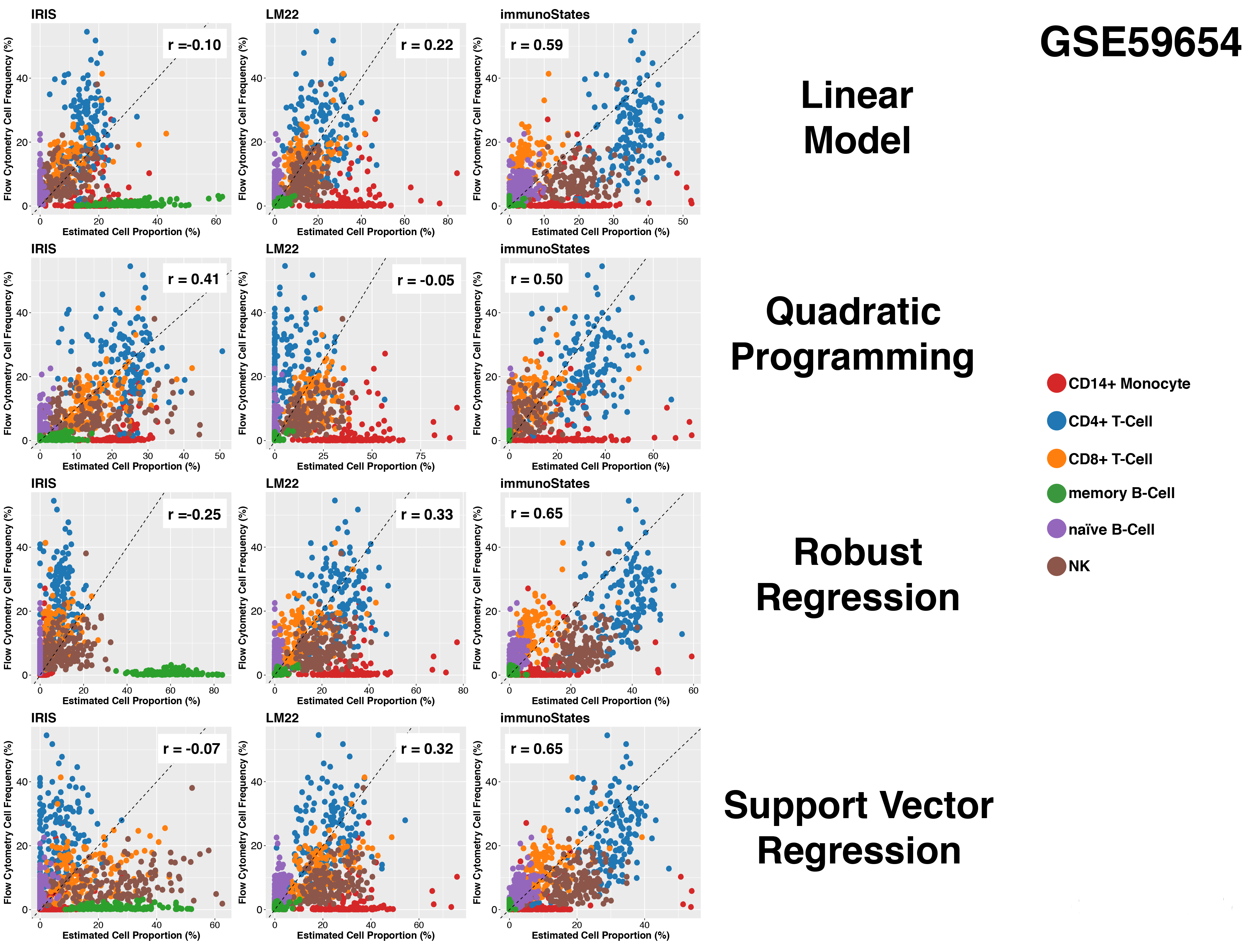
Correlation plots between measured and estimated proportions. **(5)** Correlation plots of estimated (x-axis) and measured cell proportions (y-axis) for each method and matrix combination for samples in GSE65133. Correlation is measured by Pearson’s correlation coefficient. **(6)** Same as in **(5)** for GSE59654. **(7,8,9)** Same as in **(5)** for Stanford-Ellison 2011, 2012, and 2013 sample cohorts respectively.

**Supplementary Figure 7:**
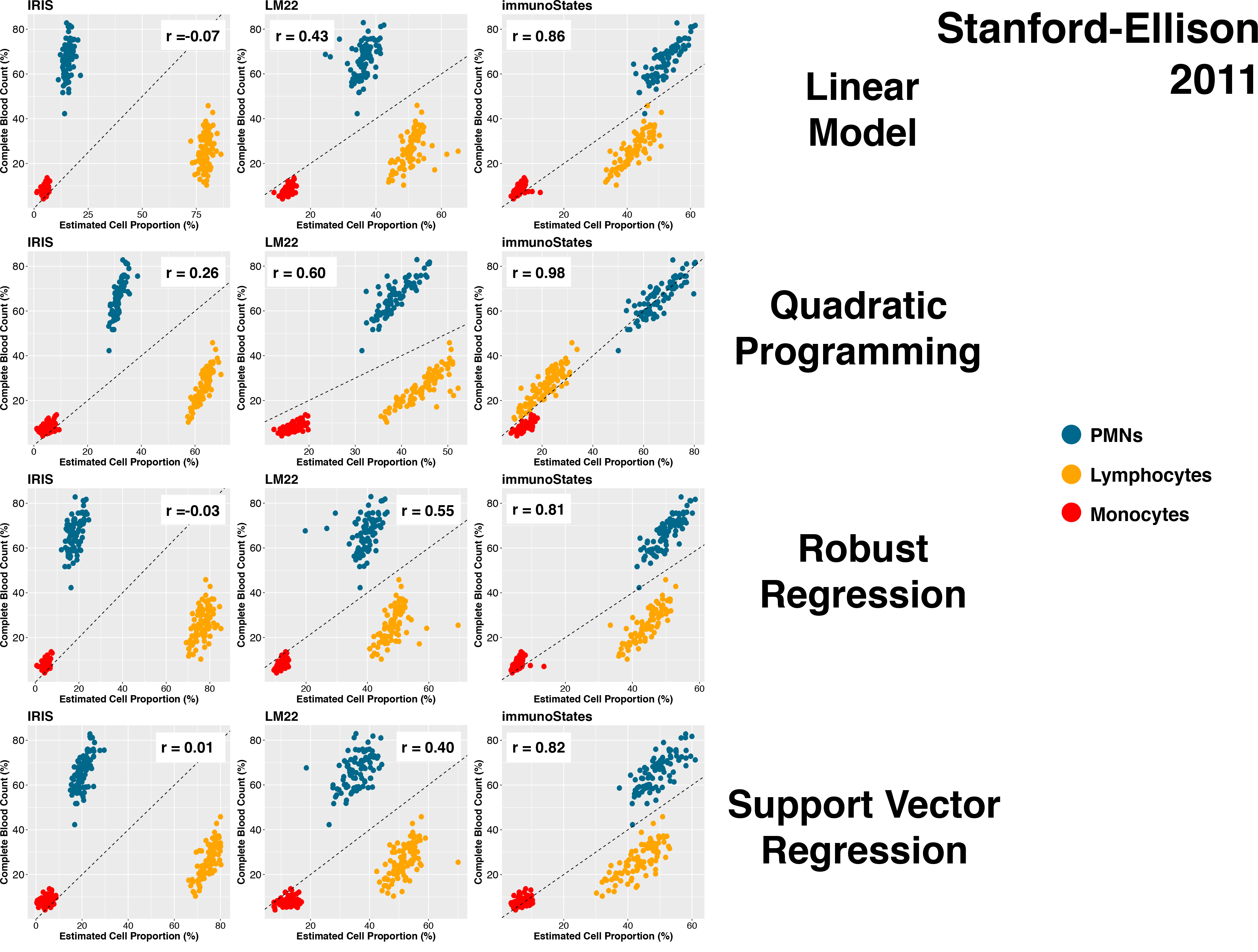
Correlation plots between measured and estimated proportions. **(5)** Correlation plots of estimated (x-axis) and measured cell proportions (y-axis) for each method and matrix combination for samples in GSE65133. Correlation is measured by Pearson’s correlation coefficient. **(6)** Same as in **(5)** for GSE59654. **(7,8,9)** Same as in **(5)** for Stanford-Ellison 2011, 2012, and 2013 sample cohorts respectively.

**Supplementary Figure 8:**
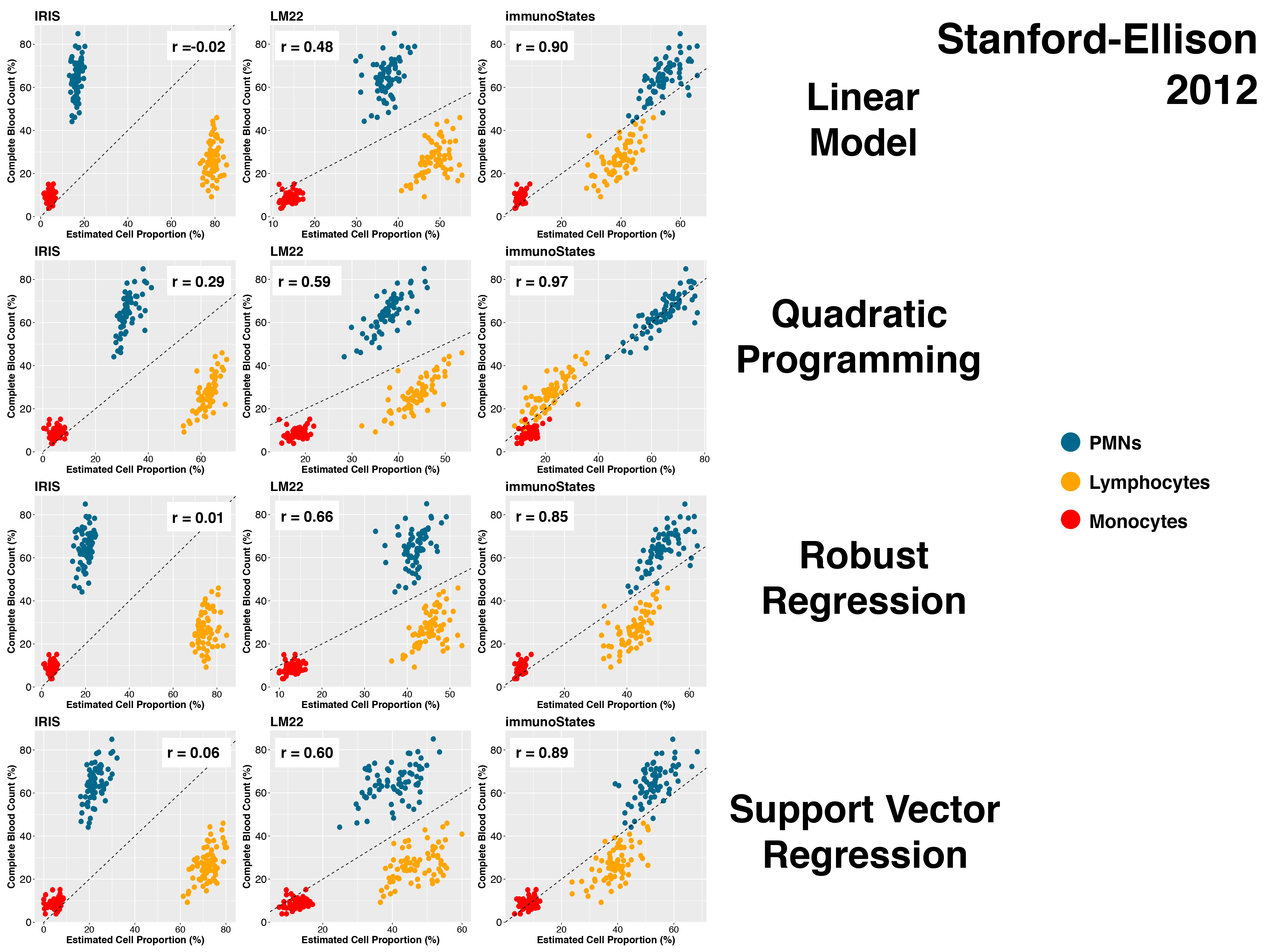
Correlation plots between measured and estimated proportions. **(5)** Correlation plots of estimated (x-axis) and measured cell proportions (y-axis) for each method and matrix combination for samples in GSE65133. Correlation is measured by Pearson’s correlation coefficient. **(6)** Same as in **(5)** for GSE59654. **(7,8,9)** Same as in **(5)** for Stanford-Ellison 2011, 2012, and 2013 sample cohorts respectively.

**Supplementary Figure 9:**
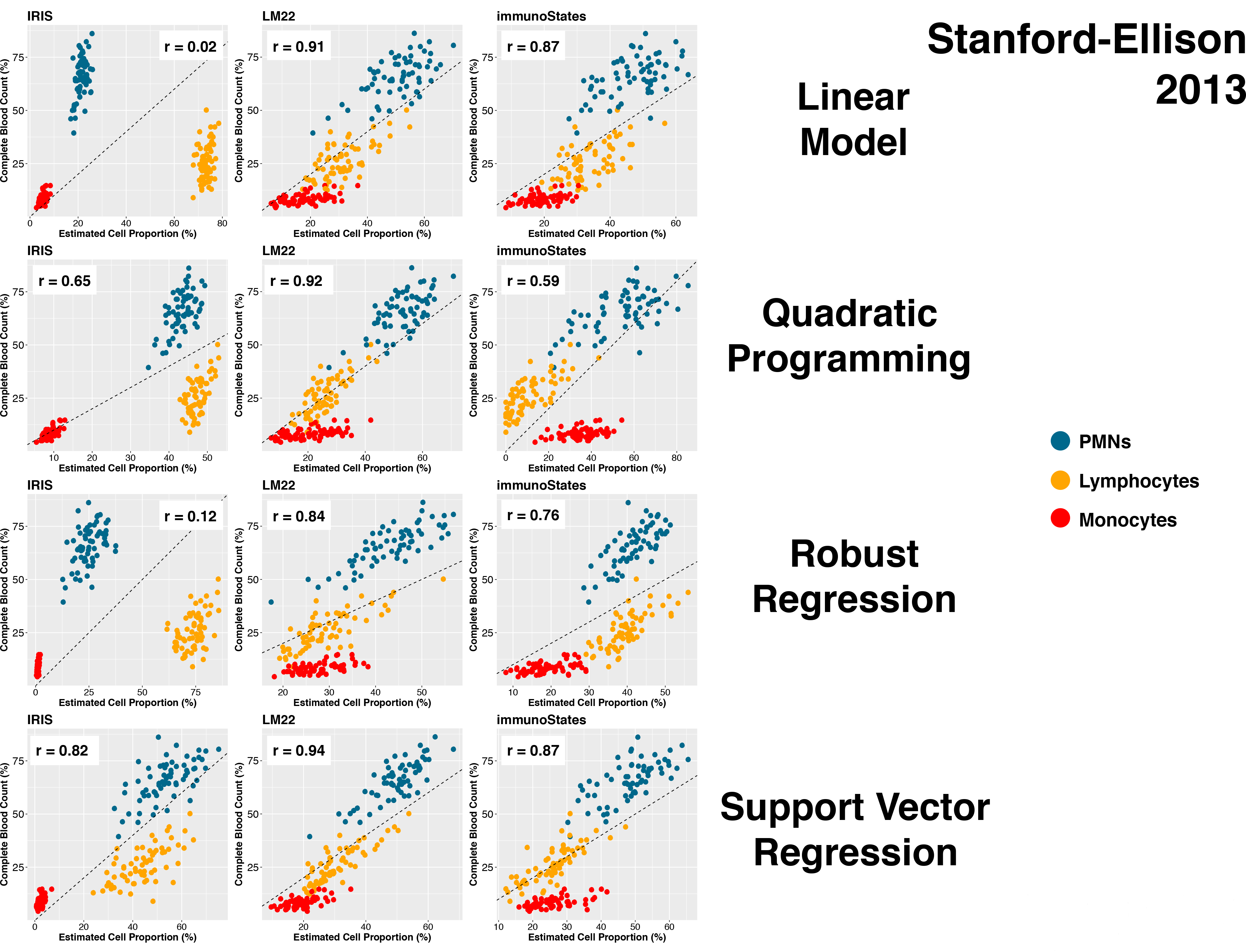
Correlation plots between measured and estimated proportions. **(5)** Correlation plots of estimated (x-axis) and measured cell proportions (y-axis) for each method and matrix combination for samples in GSE65133. Correlation is measured by Pearson’s correlation coefficient. **(6)** Same as in **(5)** for GSE59654. **(7,8,9)** Same as in **(5)** for Stanford-Ellison 2011, 2012, and 2013 sample cohorts respectively.

**Supplementary Figure 10:**
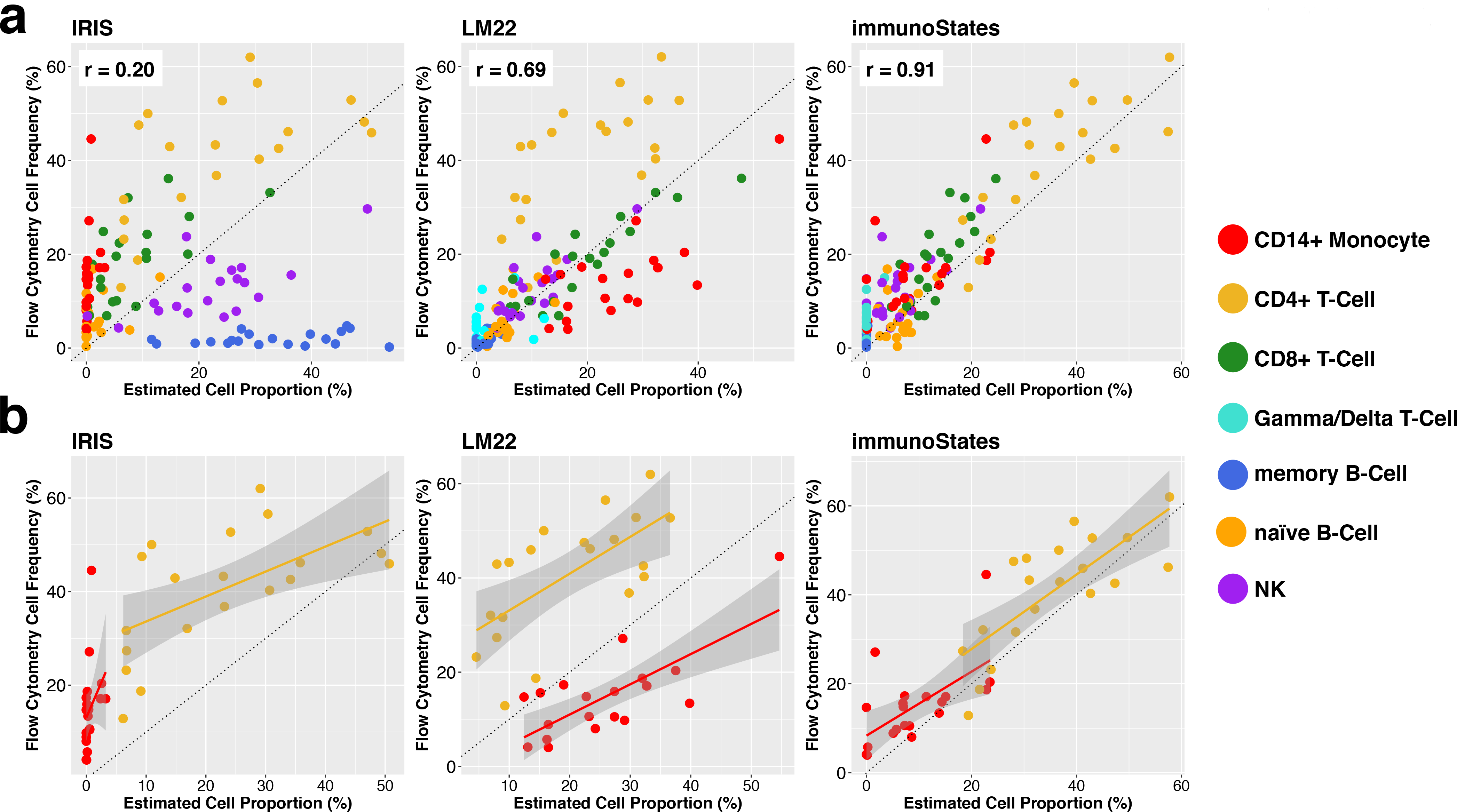
Example of systematic under-and over-estimation of cell proportions in cell mixture deconvolution. **(a)** Correlation plots of estimated (x-axis) and measured cell proportions (y-axis) in GSE65133 for all matrices using Support Vector Regression. **(b)** Highlighted CD4+ T-cell (yellow) and Monocyte (red) cell proportion estimates against measured values. Solid lines represent the best fit with a linear model, whereas the dashed line represent the 45-degree diagonal.

